# A note on predator-prey dynamics in radiocarbon datasets

**DOI:** 10.1101/2023.11.12.566733

**Authors:** Nimrod Marom, Uri Wolkowski

## Abstract

Predator-prey interactions have been a central theme in population ecology for the past century, but real-world data sets only exist for recent, relatively short (<100 years) time spans. This limits our ability to study centennial/millennial-scale predator-prey dynamics. We propose that regional radiocarbon databases can be used to reconstruct a signal of predator-prey population dynamics in deep time, overcoming this limitation. We support our argument with examples from Pleistocene Beringia and the Holocene Judean Desert.

## Introduction

Predator-prey interactions are a fundamental topic in theoretical ecology (May, 2001 [1974]), nature conservation (Johnson et al., 2019; Southall et al., 2019), and economics (Apedaille et al., 1994; Edwards et al., 2020). The dynamics of predator-prey populations exhibit multiple wavelengths, beyond the generational oscillations predicted by Lotka-Volterra models. For example, the well-known Canada lynx–snowshoe hare system oscillates on a decadal timescale, which may be linked to climatic processes operating on centennial to millennial scales (Hone et al., 2011; Yan et al., 2013). Similar multiple-timescale oscillations have been observed in other systems and predicted theoretically (Laan and Hogeweg, 1995). These oscillations may reflect multi-generational evolutionary processes.

Long-term predator-prey dynamics are difficult to study due to the scarcity of population data on timescales beyond a century. The longest record known to us is the hare-lynx records of the Hudson Bay Company, which reflect a century of fur trade (Elton and Nicholson, 1942). Other records are much shorter, typically covering decades (Gilg et al., 2009; e.g., Vucetich et al., 2011). We propose that regional sets of radiocarbon-dated animal remains can be used to study predator-prey dynamics in deep time. Because radiocarbon can date materials up to 50,000 years old, it can extend the timescale for studying these important ecological interactions by three orders of magnitude.

Radiocarbon dating is best known for providing absolute dates for archaeological and paleontological organic materials, anchoring stratigraphic sequences and establishing the temporal context of specific findings. Large radiocarbon databases are also used in archaeology to infer changes in human demography (e.g., Stewart et al., 2021) or mammalian community structure (Lazagabaster et al., 2022). These demographic inferences are based on the “dates as data’’ paradigm, which assumes that the number of radiocarbon dates in a region reflects the magnitude of occupation or the total number of person-years of human existence (Rick, 1987). This is routinely applied today using summed probability distribution (SPDs), which are applied to calibrated radiocarbon dates (Williams, 2012)In paleontology, the probability of a specimen surviving to be dated is assumed to be proportional to the number of individuals of its taxon that existed in a specific region and time (Lazagabaster et al., 2022; Stuart and Lister, 2014). For example, radiocarbon data have been used to study megafaunal extinctions (Broughton and Weitzel, 2018; e.g., Stewart et al., 2021), using archaeological and paleoenvironmental data to assess the relative importance of anthropogenic and paleoclimatic drivers. Here, we address the more general question of whether the density of radiocarbon dates obtained from a regional set of paleozoological survey data can reveal long-term predator-prey population dynamics.

Radiocarbon data are inherently sparse, prone to selection and preservation biases, and subject to uncertainties arising from measurement error and calibration procedures used to adjust observed isotopic ratios to ancient background levels (Carleton, 2021; Hajdas et al., 2021; Reimer et al., 2020). Therefore, any attempt to infer ecological processes from radiocarbon dates should exercise caution, employing minimalist hypotheses, spatially constrained samples, and randomly collected specimens to minimize biases. Radiocarbon datasets spanning a wide time range with continuous deposition and multiple species occurrences are relatively resilient to sample size and effect size issues, and the number of dates in each SPD (sensu Williams, 2012)becomes less critical when identifying strong signals in long-term trends (Crema, 2022; Hinz, 2020). In addition, our method is grounded on comparing the signal of specific taxa in constrained geographical regions through time, and, providing a strong signal, our main concern should be an equivalence of sample sizes between the compare SPDs.

We found two datasets that meet the above criteria. The first comprises published radiocarbon dates of Late Pleistocene mammalian megafauna recovered from gravels near Fairbanks, Alaska (Fox-Dobbs et al., 2008; Leonard et al., 2007). The Fairbanks data includes 33 wolves (*Canis lupus*), 28 horses (*Equus* sp.), and 3 reindeer (*Rangifer tarandus*), representing regional mortality between ∼40-7 kya. The second dataset is from the Holocene (∼10-0.5 kya) southern Judean Desert, Israel, where Lazagabaster et al. (2022) collected radiocarbon dates of leopard (*Panthera pardus nimr*, N = 12), hyrax (*Procavia capensis*, N = 27), and Nubian ibex (*Capra ibex nubiana*, N = 10) from biogenic cave deposits. The Judean Desert data are argued to represent a random sample of the regional fauna (Lazagabaster et al., 2022).

We hypothesize that the summed probability distribution (SPD) of predator radiocarbon dates, insofar as it tracks changes in population size, will have either greater or lesser divergence than expected from a random SPDs sampling in a homogeneous distribution from the same time range. A non-random divergence would suggest that predator and prey populations covaried. This minimalist hypothesis assumes nothing about the wavelength, mechanism, or cause of predator-prey interaction, which we believe cannot be tested with the current data. If supported, this hypothesis would provide preliminary evidence that long-term regional radiocarbon data encode predator-prey interaction signals. This could justify constructing larger datasets to enable in-depth investigation of the structure of these signals.

It is important to emphasize that although we aim to reconstruct long-term ecological interactions between predator and prey taxa, our primary data are the distributions of observations of single taxa over time, aggregated across several find spots in each region. Therefore, biotic and abiotic biases in specimen frequencies should not affect our results unless we have reason to believe that these processes acted differently through time on specific taxa. For example, the hypothetical fact that predator tibiae preserve less well than herbivore tibiae does not matter to the distribution of predator remains over time. Conversely, if we have reason to believe that predator tibiae preserve less well during a particular time interval compared to other periods, this could bias our results. Here, we make the uniformitarian assumption that there are no changes over time in the biotic or abiotic factors affecting the deposition or post-depositional survivability of specific taxa (top predators/larger herbivores) and that the population density of a species in a region is the main factor affecting the probability of finding their remains in the paleontological record and obtaining radiocarbon dates for them. This assumption relies on the comparability of the depositional environments for each dataset throughout time, which consist of dry desert caves or gravel deposits.

In the same vein, the overrepresentation of carnivores in both datasets in relation to a real ecosystem should not affect our analysis. The estimate of predator and prey frequencies at any point in time is not obtained from their numerical ratio, which would then indeed have to reflect a reasonable predator/prey balance. Rather, it is derived from the independent calculation of the probability densities of the radiocarbon dates of each group. In this case, the frequencies are irrelevant unless they are too few to represent the distribution of the species through time, a subject to which we referred above (Crema, 2022; Hinz, 2020).

## Methods

The radiocarbon datasets, as detailed in Supplementary tables 1 and 2, were categorized into two groups for each region: predators (*Canis* / *Panthera*) and prey (*Equus, Rangifer* / *Procavia, Capra*). These groups exclude taxa that are not likely to be in trophic interaction in the Judean Desert, for example the carrion-eating striped hyena (Hyaena hyaena) or the plateau-dwelling Dorcas gazelle (Gazella dorcas). Specimens without both minimum and maximum age estimates were also excluded. The original publications by Leonard et al. (2007), Fox-Dobbs et al. (2008) and Lazagabaster et al. (2022) provide comprehensive information on the context and laboratory procedures, which are not reiterated here.

Note, however, that the leopard specimens from the Judean Desert represent bones recovered from three caves, and their estimated minimum number of individuals is six (Lazagabaster et al., 2022). Unfortunately, we cannot distinguish individuals within each cave by removing from our analysis specimens that have similar radiocarbon dates, based, e.g., on the overlap of their 95% highest posterior density. This type of ‘chronological minimum number of individuals’ calculation is conceptually equivalent to flattening the summed probability density curves we are comparing and making them ipso facto like random noise. In this analysis, therefore, we assume that the specimens are independent from each other. This is partially supported by the fact that three leopard specimens that were recovered from the same cave and that yielded aDNA belong to three different individuals, although their minimum number of individuals would be calculated as one (Davidovich et al. under revision). Regardless, we acknowledge that some degree of interdependence is expected under these conditions and cannot be controlled.

The grouped radiocarbon ages underwent calibration (using the ‘rcarbon::calibrate’ function) and were subsequently converted to summed probability distributions (SPDs) using the ‘rcarbon::spd’ function. These operations were performed in R 4.3.0 (R Core Team, 2021), utilizing the ‘rcarbon’ library developed by Crema and Bevan (2021).

Kullback-Leibler (KL) divergence (Kullback & Leibler, 1951) quantifies the distance between two probability distributions by calculating the difference between the Shannon entropy of the first distribution and the cross-entropy of the first and second distributions. The resulting KL divergence value is not a distance metric as it does not satisfy the triangle inequality and is asymmetric, meaning the divergence of *p(x)* from *q(x)* differs from the divergence of *q(x)* from *p(x)*. The KL divergence is one of the most popular ways to compare probability distributions in information theory and data science, and has mathematical properties that make it uniquely suitable for measuring relative information (reviewed in Deng et al., 2019). KL divergence from the prey to the predator SPD was computed in each case using the ‘philentropy::KL’ function from the ‘philentropy’ library (Drost, 2018). Following this, a random set of integers, equivalent to the sample size of the predator, was drawn from the range of the radiocarbon years of the prey. Each integer in the random set was assigned a radiocarbon measurement error that closely matched the real radiocarbon error in the actual dataset. These integers were then calibrated, converted to an SPD, and the Kullback-Leibler divergence from the actual prey SPD was calculated.

This random sampling procedure was repeated 100 times with replacement. The percentage of random samples was then used to estimate the likelihood of the divergence of the predator from the prey SPDs being derived from a random dataset, giving the probability of KL(Predator||Prey) ∉KL(Random||Prey). Note that bootstrap support is a conservative estimate of accuracy in most cases and should not be understood as a statistical p-value without additional, case-specific research (Hilis & Bull, 1993). The smoothed SPDs (calculated using ‘modelbased::smoothing’) are presented in the figures below. The smoothing procedure was applied after the KL divergence calculations.

## Results and concluding remarks

The Fairbanks dataset shows fluctuating and alternating values of predator and prey densities over the interval between ∼45 and 7 kya (Figure 1). The Kullback-Leibler (KL) divergence between the predator and prey distributions is 1.7174, which is smaller than 98% of the divergences measured for (random) predator-(real) prey distributions. This supports our hypothesis that the Fairbanks predator and prey distributions are not random, and that the low divergence between them is therefore unlikely to be due to chance.

**Table 1.** Summary statistics of the original KL divergence of predator from prey SPD in the Fairbanks and the Judean Desert datasets.

| <u>Statistic</u> | <u>Fairbanks</u> | <u>Judean Desert</u> |
| --- | --- | --- |
| KL(Predator\|\|Prey) | 1.7174 | 5.0741 |
| KL(Random\|\|Prey) |  |  |
| Min. | 1.559 | 3.065 |
| 1st quartile | 2.148 | 4.001 |
| Median | 2.315 | 4.523 |
| Mean | 2.294 | 4.417 |
| 3rd quartile | 2.473 | 4.858 |
| Max. | 3.072 | 5.242 |
| Bootstrap support for $KL(Predator\vert \vert Prey) \notin KL(Random\vert \vert Prey)$ | 98% | 94% |

**Figure 1.**
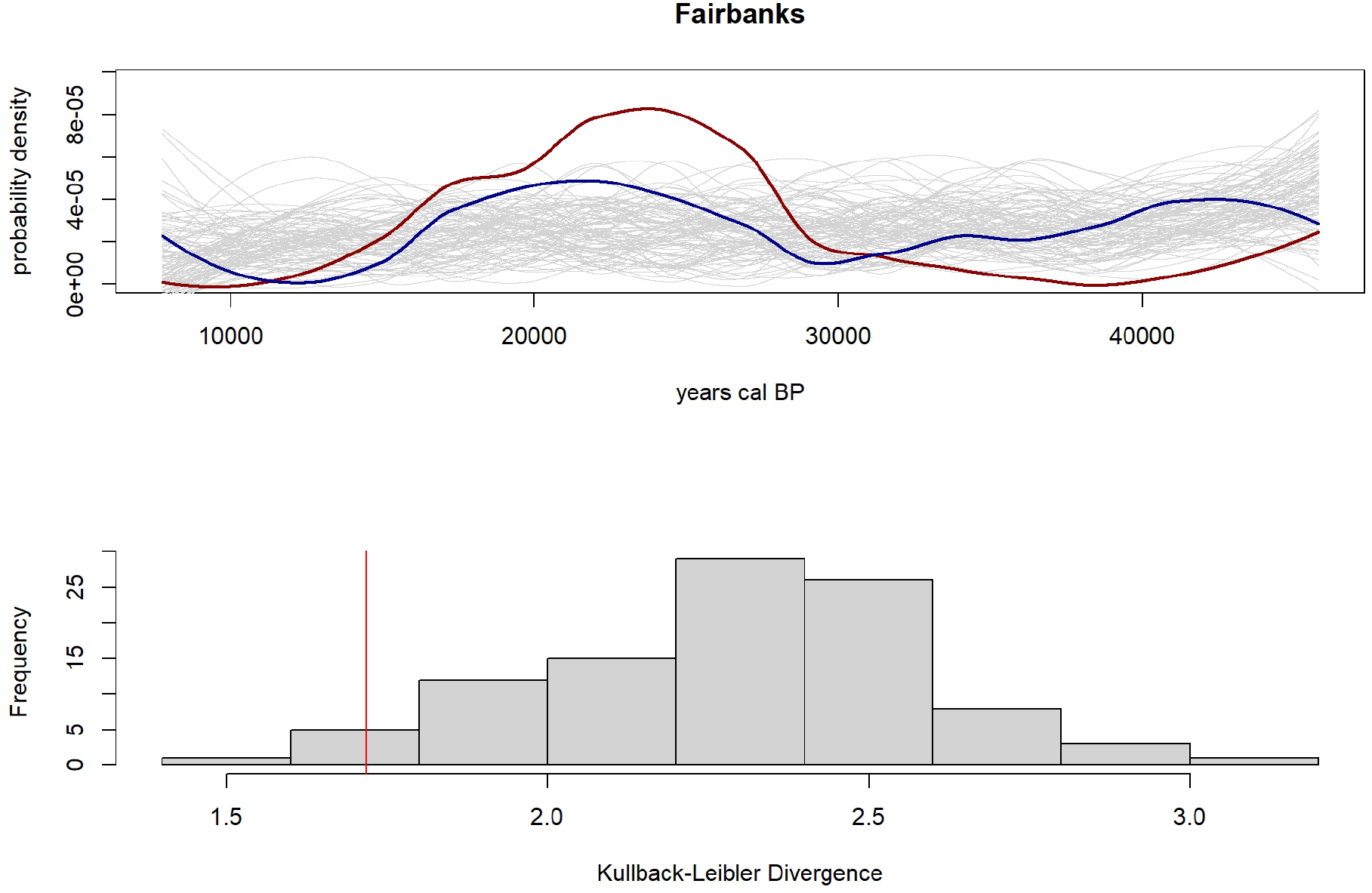
The prey (red) and predator (blue) SPD for the Fairbanks data, against the background of 100 random SPD replicates (top). The distribution of the Kullback-Leibler divergences from the prey to the predator KL(Predator||Prey) SPDs is marked by the vertical red line on the histogram below, which shows the distribution of the KL divergences between the prey SPD and the random replicates KL(Random||Prey) (bottom).^1^

Similarly, the Judean Desert dataset shows fluctuating and alternating values of predator and prey densities over the interval between 10,000 and <500 years (Figure 2). The Kullback-Leibler (KL) divergence between the predator and prey distributions is 5.0741, which is greater than 94% of the divergences measured for (random) predator-(real) prey distributions. This supports our hypothesis that the Judean Desert predator and prey distributions are not random, and that the low divergence between them is therefore unlikely to be due to chance.

**Figure 2.**
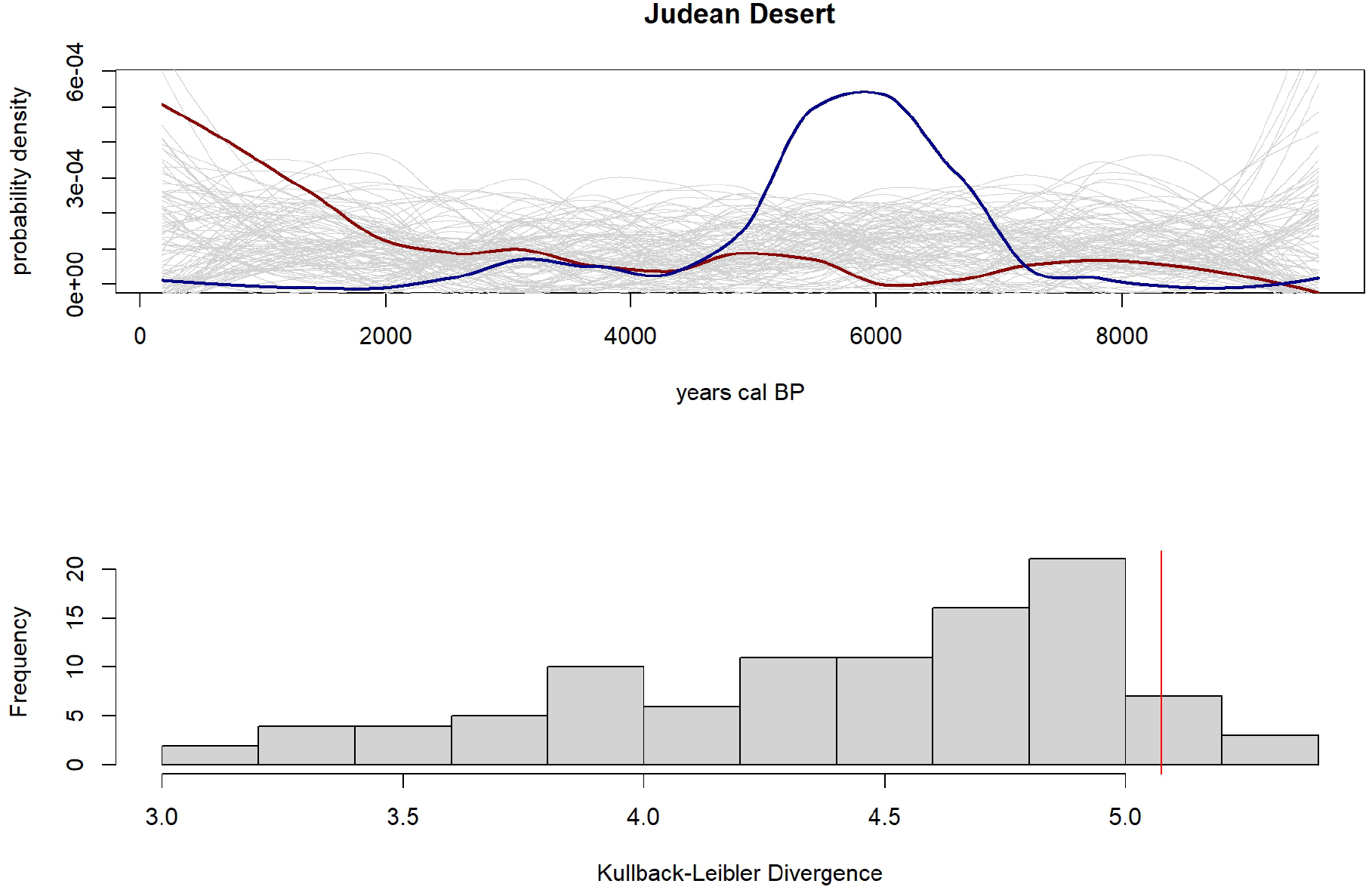
The prey (red) and predator (blue) SPD for the Judean Desert data, against the background of 100 random SPD replicates (top).^2^ The distribution of the Kullback-Leibler divergences from the prey to the predator KL(Predator||Prey) SPDs is marked by the vertical red line on the histogram below, which shows the distribution of the KL divergences between the prey SPD and the random replicates KL(Random||Prey) (bottom).

As a specific case of population dynamics, predator-prey systems are typically studied over short time periods, limiting our understanding of long-term fluctuations driven by factors such as climate change and evolution. Radiocarbon records may capture signals of these dynamics under rare sampling conditions. Here, we tested the hypothesis that the divergence between predator and prey probability density curves is not random (KL(Predator||Prey) ∉KL(Random||Prey)) using two coupled datasets from Fairbanks, Alaska, and the southern Judean Desert, Israel. Our results suggest that in these cases the divergence is unlikely to be random. This is informative of the idea that radiocarbon data may sequester long-term predator-prey interactions that are over and beyond the timescale observed until now. Additional high-resolution datasets are required to validate these results and further investigate the observed patterns.

## Appendices

**Supplementary table 1.**
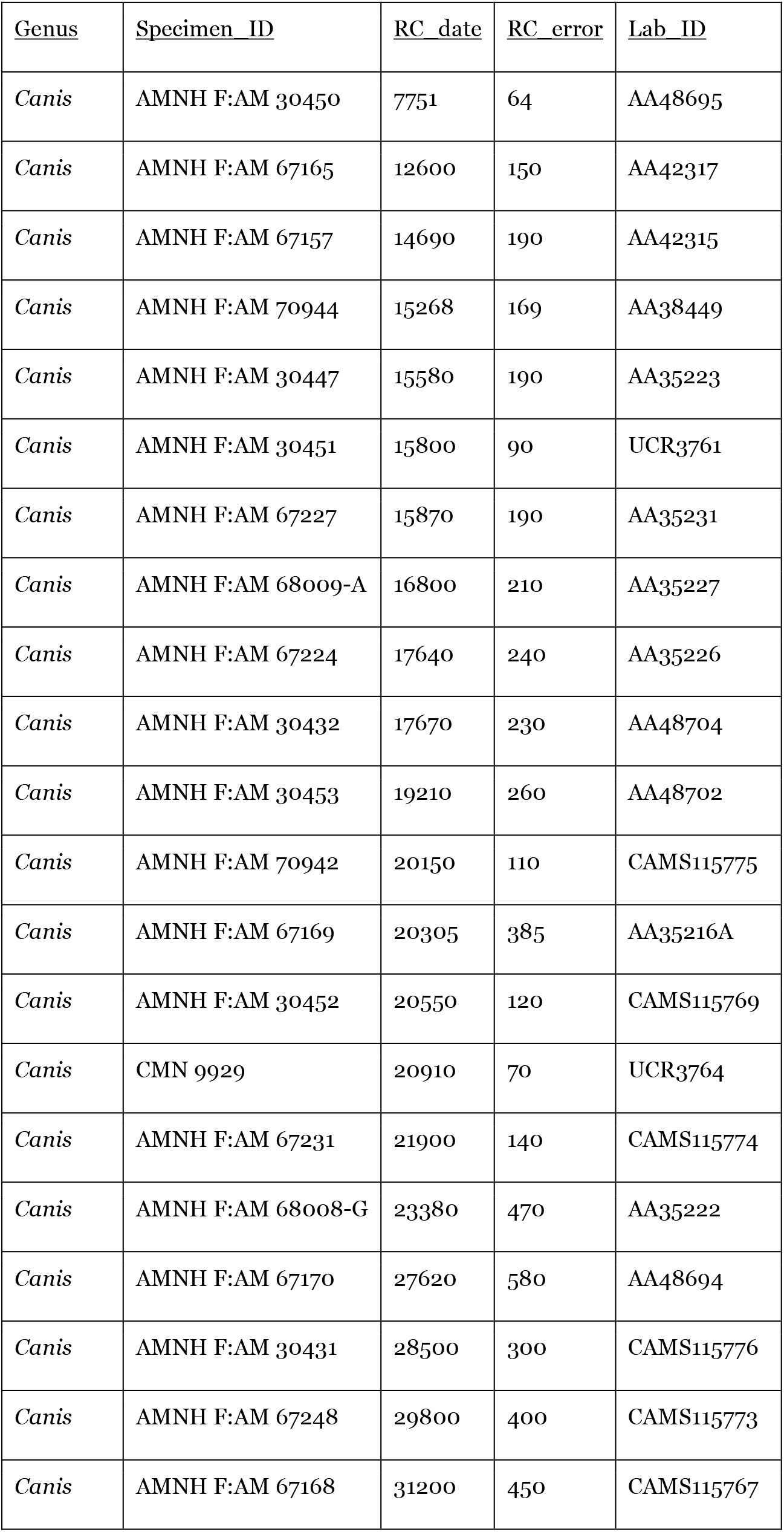
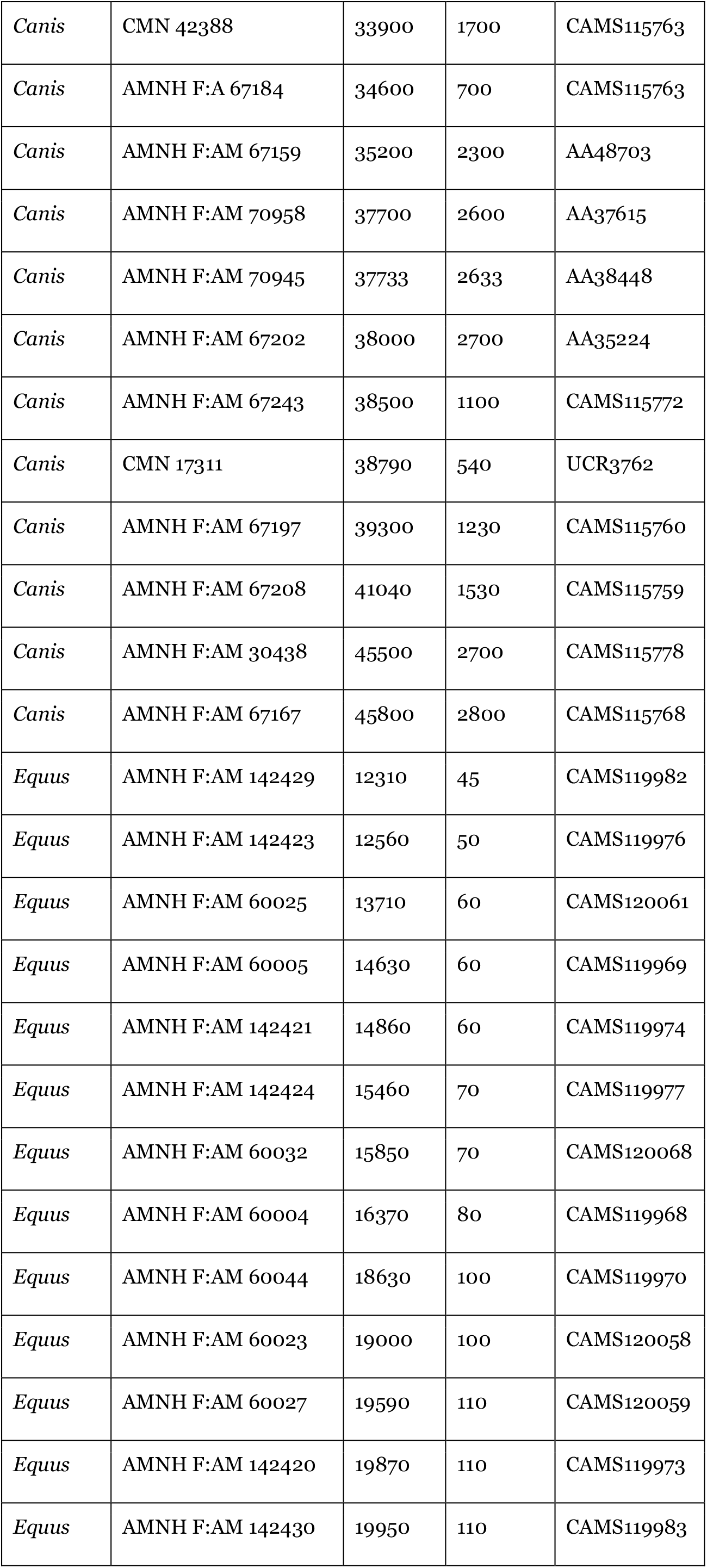
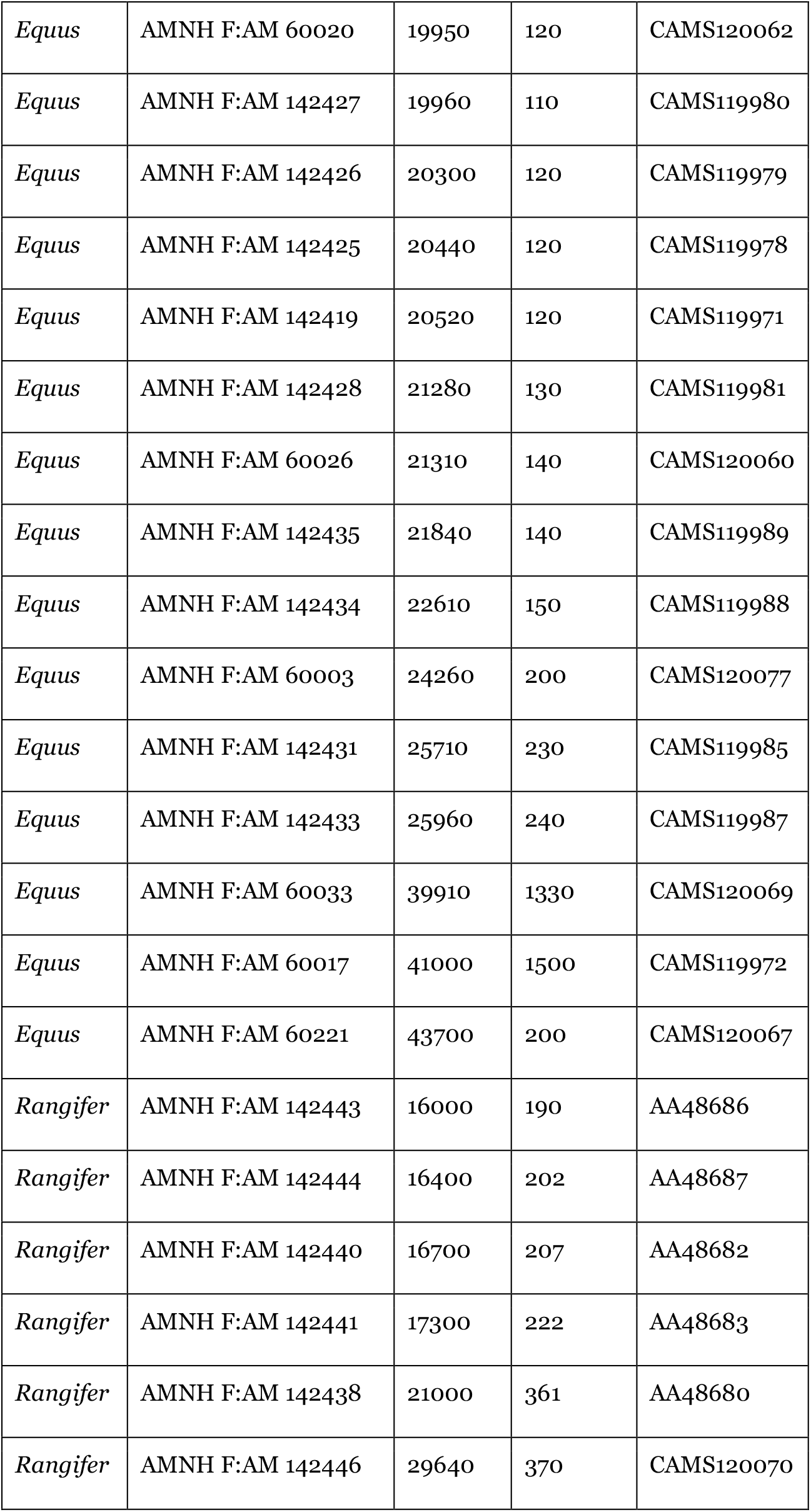
The Beringia dataset, from Fox-Dobbs et al. (2008, tables 2-3)

| <u>Genus</u> | <u>Specimen ID</u> | <u>RC date</u> | <u>RC error</u> | <u>Lab ID</u> |
| --- | --- | --- | --- | --- |
| *Canis* | AMNH F:AM 30450 | 7751 | 64 | AA48695 |
| *Canis* | AMNH F:AM 67165 | 12600 | 150 | AA42317 |
| *Canis* | AMNH F:AM 67157 | 14690 | 190 | AA42315 |
| *Canis* | AMNH F:AM 70944 | 15268 | 169 | AA38449 |
| *Canis* | AMNH F:AM 30447 | 15580 | 190 | AA35223 |
| *Canis* | AMNH F:AM 30451 | 15800 | 90 | UCR3761 |
| *Canis* | AMNH F:AM 67227 | 15870 | 190 | AA35231 |
| *Canis* | AMNH F:AM 68009-A | 16800 | 210 | AA35227 |
| *Canis* | AMNH F:AM 67224 | 17640 | 240 | AA35226 |
| *Canis* | AMNH F:AM 30432 | 17670 | 230 | AA48704 |
| *Canis* | AMNH F:AM 30453 | 19210 | 260 | AA48702 |
| *Canis* | AMNH F:AM 70942 | 20150 | 110 | CAMS115775 |
| *Canis* | AMNH F:AM 67169 | 20305 | 385 | AA35216A |
| *Canis* | AMNH F:AM 30452 | 20550 | 120 | CAMS115769 |
| *Canis* | CMN 9929 | 20910 | 70 | UCR3764 |
| *Canis* | AMNH F:AM 67231 | 21900 | 140 | CAMS115774 |
| *Canis* | AMNH F:AM 68008-G | 23380 | 470 | AA35222 |
| *Canis* | AMNH F:AM 67170 | 27620 | 580 | AA48694 |
| *Canis* | AMNH F:AM 30431 | 28500 | 300 | CAMS115776 |
| *Canis* | AMNH F:AM 67248 | 29800 | 400 | CAMS115773 |
| *Canis* | AMNH F:AM 67168 | 31200 | 450 | CAMS115767 |
| *Canis* | CMN 42388 | 33900 | 1700 | CAMS115763 |
| *Canis* | AMNH F:A 67184 | 34600 | 700 | CAMS115763 |
| *Canis* | AMNH F:AM 67159 | 35200 | 2300 | AA48703 |
| *Canis* | AMNH F:AM 70958 | 37700 | 2600 | AA37615 |
| *Canis* | AMNH F:AM 70945 | 37733 | 2633 | AA38448 |
| *Canis* | AMNH F:AM 67202 | 38000 | 2700 | AA35224 |
| *Canis* | AMNH F:AM 67243 | 38500 | 1100 | CAMS115772 |
| *Canis* | CMN 17311 | 38790 | 540 | UCR3762 |
| *Canis* | AMNH F:AM 67197 | 39300 | 1230 | CAMS115760 |
| *Canis* | AMNH F:AM 67208 | 41040 | 1530 | CAMS115759 |
| *Canis* | AMNH F:AM 30438 | 45500 | 2700 | CAMS115778 |
| *Canis* | AMNH F:AM 67167 | 45800 | 2800 | CAMS115768 |
| *Equus* | AMNH F:AM 142429 | 12310 | 45 | CAMS119982 |
| *Equus* | AMNH F:AM 142423 | 12560 | 50 | CAMS119976 |
| *Equus* | AMNH F:AM 60025 | 13710 | 60 | CAMS120061 |
| *Equus* | AMNH F:AM 60005 | 14630 | 60 | CAMS119969 |
| *Equus* | AMNH F:AM 142421 | 14860 | 60 | CAMS119974 |
| *Equus* | AMNH F:AM 142424 | 15460 | 70 | CAMS119977 |
| *Equus* | AMNH F:AM 60032 | 15850 | 70 | CAMS120068 |
| *Equus* | AMNH F:AM 60004 | 16370 | 80 | CAMS119968 |
| *Equus* | AMNH F:AM 60044 | 18630 | 100 | CAMS119970 |
| *Equus* | AMNH F:AM 60023 | 19000 | 100 | CAMS120058 |
| *Equus* | AMNH F:AM 60027 | 19590 | 110 | CAMS120059 |
| *Equus* | AMNH F:AM 142420 | 19870 | 110 | CAMS119973 |
| *Equus* | AMNH F:AM 142430 | 19950 | 110 | CAMS119983 |
| *Equus* | AMNH F:AM 60020 | 19950 | 120 | CAMS120062 |
| *Equus* | AMNH F:AM 142427 | 19960 | 110 | CAMS119980 |
| *Equus* | AMNH F:AM 142426 | 20300 | 120 | CAMS119979 |
| *Equus* | AMNH F:AM 142425 | 20440 | 120 | CAMS119978 |
| *Equus* | AMNH F:AM 142419 | 20520 | 120 | CAMS119971 |
| *Equus* | AMNH F:AM 142428 | 21280 | 130 | CAMS119981 |
| *Equus* | AMNH F:AM 60026 | 21310 | 140 | CAMS120060 |
| *Equus* | AMNH F:AM 142435 | 21840 | 140 | CAMS119989 |
| *Equus* | AMNH F:AM 142434 | 22610 | 150 | CAMS119988 |
| *Equus* | AMNH F:AM 60003 | 24260 | 200 | CAMS120077 |
| *Equus* | AMNH F:AM 142431 | 25710 | 230 | CAMS119985 |
| *Equus* | AMNH F:AM 142433 | 25960 | 240 | CAMS119987 |
| *Equus* | AMNH F:AM 60033 | 39910 | 1330 | CAMS120069 |
| *Equus* | AMNH F:AM 60017 | 41000 | 1500 | CAMS119972 |
| *Equus* | AMNH F:AM 60221 | 43700 | 200 | CAMS120067 |
| *Rangifer* | AMNH F:AM 142443 | 16000 | 190 | AA48686 |
| *Rangifer* | AMNH F:AM 142444 | 16400 | 202 | AA48687 |
| *Rangifer* | AMNH F:AM 142440 | 16700 | 207 | AA48682 |
| *Rangifer* | AMNH F:AM 142441 | 17300 | 222 | AA48683 |
| *Rangifer* | AMNH F:AM 142438 | 21000 | 361 | AA48680 |
| *Rangifer* | AMNH F:AM 142446 | 29640 | 370 | CAMS120070 |

**Supplementary table 2.** The Judean Desert Holocene dataset, modified from Lazagabaster et al. (2022).

| Genus | Specimen_ID | Material | RCdate | RCerror | Laboratory | Lab_ID |
| --- | --- | --- | --- | --- | --- | --- |
| Panthera | HE-004 | collagen | 5317 | 25 | OxA | 39091 |
| Panthera | HE-008 | collagen | 5301 | 25 | OxA | 39214 |
| Panthera | HE-010 | collagen | 4477 | 26 | OxA | 38732 |
| Panthera | HE-053 | collagen | 4508 | 30 | OxA | 38788 |
| Panthera | HE-054 | collagen | 6239 | 26 | OxA | 39216 |
| Panthera | HE-124 | collagen | 5375 | 28 | OxA | 47465 |
| Panthera | HE-195 | collagen | 4945 | 25 | OxA | 39107 |
| Panthera | HE-230 | collagen | 4935 | 26 | OxA | 39109 |
| Panthera | HE-231 | collagen | 5312 | 25 | OxA | 39110 |
| Panthera | HE-232 | collagen | 5366 | 30 | OxA | 47473 |
| Panthera | SK-703 | bioapatite | 3200 | 30 | UGAMS | 44190 |
| Panthera | UR-035 | bioapatite | 8960 | 30 | UGAMS | 48266 |
| Capra | EG-002 | bioapatite | 7850 | 30 | UGAMS | 46061 |
| Capra | HE-021 | collagen | 3141 | 22 | OxA | 39215 |
| Capra | HE-024 | collagen | 1860 | 20 | UGAMS | 46066 |
| Capra | QN-028 | collagen | 1870 | 20 | OxA | 39255 |
| Capra | QN-083 | collagen | 4550 | 24 | OxA | 39175 |
| Capra | SK-014 | collagen | 350 | 20 | UGAMS | 46081 |
| Capra | TS-022 | collagen | 1338 | 20 | OxA | 39227 |
| Capra | QN-084 | collagen | 4520 | 20 | UGAMS | 46078 |
| Capra | TZB-004 | collagen | 870 | 20 | UGAMS | 48253 |
| Capra | TS-032 | collagen | 1060 | 20 | UGAMS | 46116 |
| Procavia | ARN-001 | collagen | 180 | 20 | UGAMS | 48283 |
| Procavia | ARN-002 | bioapatite | 9600 | 110 | UGAMS | 48284 |
| Procavia | C513-003 | collagen | 430 | 20 | UGAMS | 48261 |
| Procavia | C513-004 | collagen | 1490 | 20 | UGAMS | 48260 |
| Procavia | EG-009 | collagen | 7141 | 28 | OxA | -88 |
| Procavia | HE-005 | collagen | 2948 | 22 | OxA | 39213 |
| Procavia | HE-100 | collagen | 2920 | 20 | UGAMS | 46068 |
| Procavia | HE-153 | collagen | 6476 | 26 | OxA | 39218 |
| Procavia | HE-177 | bioapatite | 4350 | 20 | UGAMS | 46069 |
| Procavia | HE-184 | collagen | 2920 | 22 | OxA | 39106 |
| Procavia | HYS-002 | collagen | 1130 | 20 | UGAMS | 49003 |
| Procavia | HYS-003 | collagen | 1280 | 20 | UGAMS | 49004 |
| Procavia | QN-026 | collagen | 924 | 19 | OxA | 39255 |
| Procavia | QN-038 | collagen | 2520 | 30 | UGAMS | 46077 |
| Procavia | QN-056 | collagen | 1144 | 20 | OxA | 39260 |
| Procavia | QN-087 | collagen | 335 | 20 | OxA | 39176 |
| Procavia | SK-001 | collagen | 363 | 22 | OxA | 39607 |
| Procavia | SK-002 | collagen | 376 | 19 | OxA | 39608 |
| Procavia | SK-046 | collagen | 1590 | 20 | OxA | 39609 |
| Procavia | SK-152 | bioapatite | 6490 | 30 | UGAMS | 46134 |
| Procavia | SK-189 | collagen | 1140 | 20 | UGAMS | 46085 |
| Procavia | SK-211 | collagen | 4228 | 24 | OxA | 39930 |
| Procavia | SK-241 | collagen | 620 | 18 | OxA | 39615 |
| Procavia | SK-493 | collagen | 646 | 18 | OxA | 39849 |
| Procavia | TS-065 | collagen | 1160 | 19 | OxA | 47371 |
| Procavia | TZB-009 | collagen | 2010 | 20 | UGAMS | 48251 |
| Procavia | YO-013 | collagen | 721 | 18 | OxA | 39851 |

## Acknowledgements

We thank Jesús Rodríguez, Miriam Belmaker, Shimon Maromand an anonymous reviewer for their comments on the manuscript, and to Ruth Blasco for recommending it.

## Data, scripts, code, and supplementary information availability

Data and code are available online: https://zenodo.org/doi/10.5281/zenodo.10115388

## Footnotes

1 The function was run with the following parameters: kld_dates(fairbanks_prey$RC_date, fairbanks_prey$RC_error, fairbanks_predator$RC_date, fairbanks_predator$RC_error, sample = 100, dataset_name = “Fairbanks”, y_scaling_parameter = 4, smoothing = 0.4)

2 The function was called with the following parameters: kld_dates(desco_prey$RC_date, desco_prey$RC_error, desco_predator$RC_date, desco_predator$RC_error, sample = 100, dataset_name = “Judean Desert”, y_scaling_parameter = 4, smoothing = 0.4).

